# Inoculation with *Pseudomonas fluorescens* UM270 alters the maize root-associated endobiome and interacting networks in a milpa model

**DOI:** 10.1101/2023.05.15.540877

**Authors:** Blanca Rojas-Sánchez, Hugo Castelán-Sánchez, Gustavo Santoyo

## Abstract

The traditional milpa system is a polyculture originating in Mesoamerica, whose core is maize (*Zea mays* L.), associated with squash (*Cucurbita* spp.) and beans (*Phaseolus vulgaris* L.). In recent years, milpa-type crops have decreased owing to climate change, rapid population growth, and the excessive use of agrochemicals; therefore, the application of plant growth-promoting rhizobacteria (PGPR) to counteract these negative effects has been little explored. In this study, a maize crop in a milpa system was fertilized with the PGPR *Pseudomonas fluorescens* UM270, and the endophytic root microbiome (endobiome) of maize was assessed by 16S rRNA and internal transcribed spacer regions (ITS) sequencing. The results showed that UM270 the rhizosphere inoculation of *P. fluorescens* UM270 did not increase alpha diversity in either monoculture or the milpa, but it did alter the endophytic microbiome of maize plant roots by stimulating the presence of bacterial operational taxonomic units (OTUs) of the genera *Burkholderia* and *Pseudomonas* (in a monoculture), whereas in the milpa system, the PGPR stimulated a greater endophytic diversity and the presence of genera such as *Burkholderia, Variovorax*, and N-fixing rhizobia genera, including *Rhizobium, Mesorhizobium* and *Bradyrhizobium*. No clear association was found between fungal diversity and the presence of strain UM270, but beneficial fungi such as *Rizophagus irregularis* and *Exophiala pisciphila* were detected in the milpa system. In addition, network analysis revealed unique interactions with species like *Stenotrophomonas* sp., *Burkholderia xenovorans*, and *Sphingobium yanoikuyae*, which would potentially be playing a beneficial role with the plant. To the best of our knowledge, this is the first study in which the root microbiome of maize growing under a milpa model was assessed by bio-inoculation with PGPRs.

## Introduction

Maize was one of the crops that had the greatest impact since the beginning of the green revolution, thanks to its nutritional characteristics and its high demand within the food, balanced feed, and pharmaceutical industries, and in recent years for its use in the production of bioethanol, is one of the most important cereals worldwide (Edoghogho-Imade and Olubukola-Oluranti 2021). However, its establishment as a monoculture has increased the presence and resistance of pests and diseases, and the soils are deteriorating and increasing their intoxication due to the large amounts of agrochemicals that are supplied during the development of the crop. (Camacho 2017; Jasso-Miranda et al. 2022; Prasad et al. 2018; Ureta et al. 2020).

Today, it is necessary to implement different crop systems that include the use of technologies that are environmentally friendly and counteract the effects caused by the beginning of the green revolution. (Dellepiane et al. 2015; Salazar Barrientos et al. 2016). One of the systems that has regained importance in recent years is the traditional “milpa” system, one of its characteristics is that they apply minimum or zero tillage, do not need irrigation systems, and are based on the establishment of maize cultivation associated with other crops such as beans and squash. This co-culture is known as the “TM” (TM) (Álvarez-Buylla et al. 2011; Ebel et al. 2017), where maize serves as a support for the entangling of beans, which in turn, through the production of nodules, increases nitrogen fixation that is used by maize and squash, and the latter provides soil protection by reducing the growth of weeds, and maintains the humidity and through the production of allelopathic compounds (cucurbits) released by the leaching of the rain, they keep insects away. It has been one of the most used systems over the years in Mexico, and its importance encompasses cultural, economic, social, and environmental aspects (Ku-Pech et al. 2019; Rodríguez and Arias de Reyna 2014).

On the other hand, the beneficial effects of inoculating plant growth-promoting rhizobacteria or PGPR in a milpa agrosystem, has not been explored yet. PGPRs are among the plant growth-promoting microorganisms that have been widely studied and applied in multiple agricultural crops, including maize, providing multiple services, such as the biocontrol of pathogens and of course, stimulating the plant growth and production (dos Santos et al. 2018; Molina-Romero et al. 2015; Vandana et al. 2020). Some of the most studied PGPR genera in the maize rhizosphere include *Burkholderia, Bacillus, Azotobacter, Streptomyces, Paenibacillus, Sphingobium*, and *Pseudomonas*. Pseudomonads stand out for their effectiveness as plant growth promoters in maize plants, fungicides against diseases such as *Rhizoctonia solani*, biostimulants that mitigate water stress, and bioremediators of copper toxicity in maize crops (Chu et al. 2020; Edoghogho-Imade and Olubukola-Oluranti 2021; Rana et al. 2019; Sah et al. 2017). *Pseudomonas fluorescens* strain UM270 has various PGP mechanisms, such as the production of siderophores, antibiotics, volatiles, ACC deaminase activity, biofilm formation, and phosphate solubilization. It has been proven to be an excellent promoter of plant growth *in vitro* in plants, such as *Solanum lycopersicum, Physalis ixocarpa, Medicago truncatula*, and antagonists of fungal pathogens such as *Botrytis cinerea* and *Fusarium oxysporum* (Hernández-León et al. 2015; Hernández-Salmerón et al. 2016, 2017, 2018). However, its beneficial effects in maize plants growing under a milpa model are unkown. Therefore, the objective of this study was to evaluate whether the *P. fluorescens* UM270 inoculation on maize plants stimulates the abundance of a beneficial endophytic microbiome with potential positive effects in a milpa system.

## Materials and methods

### Experimental Site

The experiment was conducted in the town of Santa Clara del Cobre in the municipality of Salvador Escalante, Michoacán. Mexico. It is located at 19° 24′ 23″ North, y 101° 38′ 24″ West at an altitude of 2239 m. The prevailing climate is humid subtropical (Köppen climate classification: Cwa). Prior to the experiment, soil analysis was carried out to determine its physicochemical characteristics. Soil analysis determined that the type of soil is clay, it is composed of a percentage of 40% sand, 41.96% clay and 18% silt, the physical and chemical characteristics of the soil are presented in Table 1.

**Table 1.**
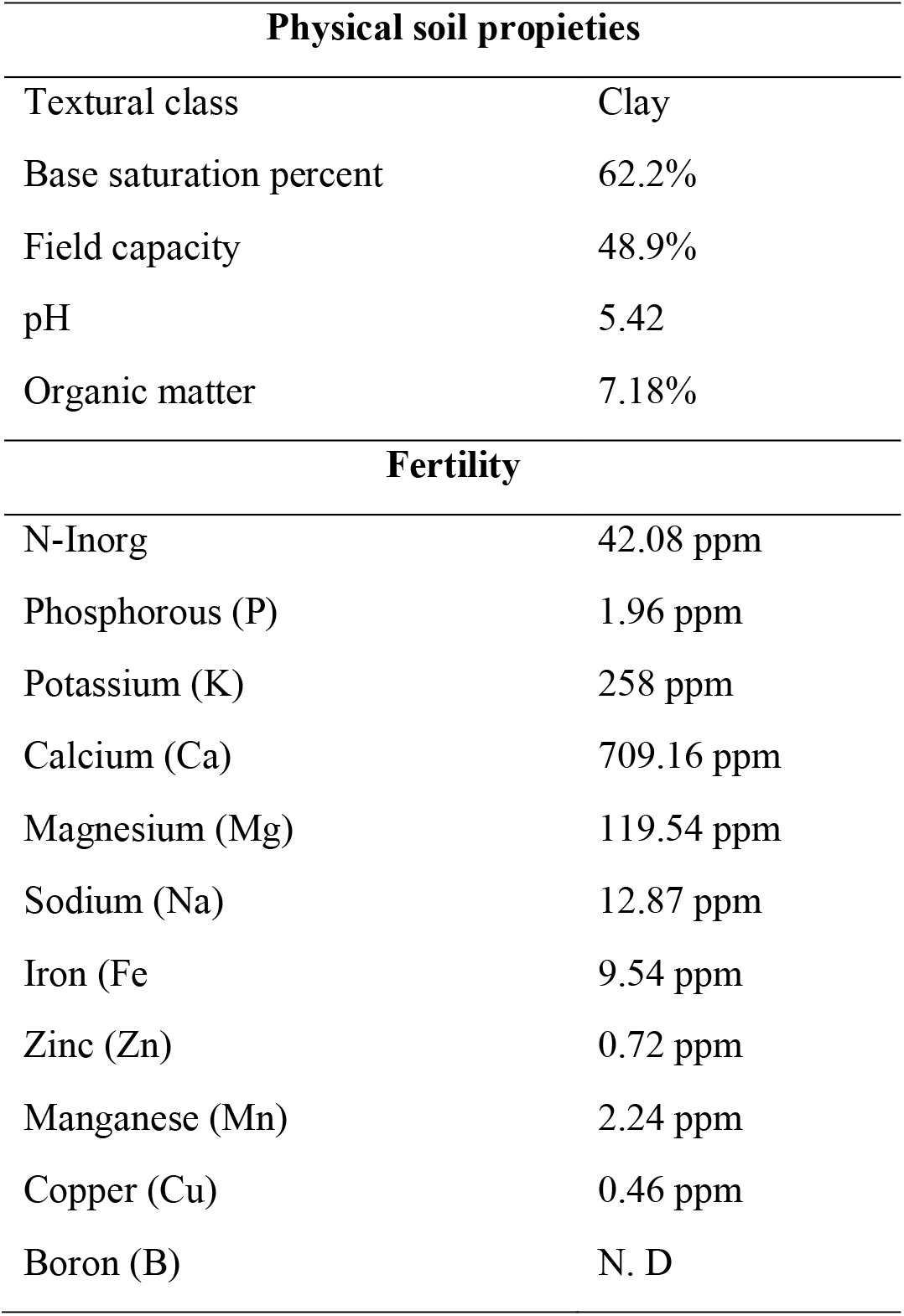
Physico-chemical characteristics of the experimental soil.

### Biological material

The seeds of *Zea mays* L., *Phaseolus vulgaris* L., and *Cucurbita* spp. used in this experiment were obtained from the same municipality where the experiment was established and were provided by the local producers. The UM270 strain was used as a bioinoculant and has been previously isolated and characterized (Hernández-León, et al., 2015).

### Inoculum preparation

Bacterial activation of *Pseudomonas fluorescens* strain UM270 was carried out by removing a hoe from the bacteria and placing it in a flask with 500 mL of Nutrient Broth (BD BIOXON), keeping it under constant agitation at 120 rpm at 28 °C for 24 h. h until an optical density (560-600 nm) of 1 was reached, and the separation of the supernatant and the bacterial pellet was carried out to subsequently suspend it in solution with 0.1 mM magnesium sulfate and finally a count of colony forming units (CFU) per milliliter.

### Seed treatments

Seed preparation consisted of a superficial disinfection process involving washing with 70% ethanol, 5% sodium hypochlorite, and sterile distilled water. The seeds used for the treatments in the presence of the bacterial strain were inoculated at a concentration of approximately 1×10^3^ CFU per seed.

### Establishment of the experiment in the field

Prior to the establishment of the crop, traditional cultural tasks were performed, including dragging, lumping, and furrowing. Maize planting was carried out on May 11, 2021 and the entire stage of cultivation ended in December of the same year, native maize seeds known as “white maize” were used. This variety is selected in the area for its characteristics of nixtamalization and tortilla flavor. After two weeks, guide beans and squash were planted. One month after planting maize, a second inoculation with the UM270 strain at a concentration of 1×10^8^ UFC was carried out on the crops with the inoculated seeds, and after another month, a third inoculation was carried out at the same concentration.

### Experimental design

The experimental design was completely randomized with three treatments, in which the three crops were planted at different planting densities. According to recommendations from the producers in the region, eight maize plants m^2^ were planted, with a n=100 plants in each treatment. The composition of the polycultures was calculated as follows: planting a maize plant is equivalent to 0.75 bean plants and 0.25 squash plants. The treatments evaluated were: (1) *Zea mays* L. (Maize roots); (2) *Zea mays* L. + UM270 (Maize roots + UM270); (3) *Zea mays* L. + UM270 + *Phaseolus vulgaris* L. + *Cucurbita* spp. (Maize roots + UM270 + Milpa system).

### DNA extraction and Illumina sequencing

Three samples composed of ten healthy maize plant roots (1 g of lateral root tissue from each plant) were pooled to isolate genomic DNA and used to sequence the endophytic microbiome, including bacteria and fungi. Briefly, soil particles were removed, and root tissues were washed and superficially sterilized by immersion in 70% ethanol for 30 s, then in a 2.5% solution of commercial bleach for 5 min, followed by at least five times washing with sterile distilled water. To further confirm the sterilization process, an aliquot from the last rinse of sterile distilled water was cultured on plates with nutrient agar medium and incubated at 28 °C for 72 h. No growth of bacterial or fungal colonies was observed in such plates after incubation.

Then, plant root tissues were macerated using mortars in liquid nitrogen under sterile conditions, following the DNA extraction protocol published by (Mahuku 2004) and further purified with a DNA purification kit (PROMEGA). The quantity and quality of DNA were confirmed by electrophoresis on agarose gels stained with GelRed and visualized under UV light using a NanoDrop 1000 spectrophotometer (Thermo Scientific, Rockford, IL, USA). Nine samples (three from each treatment) with good quantity and purity were sequenced using the Illumina MiSeq platform at Mr. DNA company (Texas, USA). DNA libraries were constructed by amplifying the V3-V4 hypervariable region of the 16S rRNA genes and ITS regions using Mr. DNA. Subsequently, these amplicons were tagged and attached to PNA PCR Clamps to reduce plastid/mitochondrial DNA amplification (Gómez-Lama et al. 2021).

### Data processing

The taxonomic levels of phyla and genera were examined and are indicated for the 16S rRNA gene and ITS sequences obtained with paired-end reads. The sequences were aligned and processed using the Parallel-META 3.5 workflow (Jing et al. 2017). Operational taxonomic unit (OTU) clustering was performed using the SILVA database integrated into Parallel-META 3.5 using a 97% homology criterion. There must be at least two sequences; the minimum zero abundance criterion is 10%, and the average abundance threshold is 0.1%. The maximum and minimum abundances were set to 0.1% and 0%, respectively (Jing et al., 2017).

### Analysis Alpha and Beta diversities

The alpha and beta diversities of the sequences were examined, and all statistical analyses were performed using the R script in Parallel-META-3.5 (do et al., 2017).

### Endobiome network analysis

Endobiome network analysis involves the construction and analysis of networks representing relationships between different species within the endobiome. For this analysis, we used the igraph library, a network analysis library for R. This library provides a wide range of tools and functions for network construction, analysis, and visualization.

### Sequence accession numbers

The raw sequences are available at NCBI under BioProject accession numbers PRJNA901513 and Sequence Read Archive (SRA) accession numbers SRR22351342, SRR22351344, SRR22351348, SRR22351343, SRR22351346, SRR22351345, SRR22351347, and SRR22351341.

### Statistical analysis

The data obtained were analyzed by analysis of variance, and the variables that presented significant differences were analyzed by Tukey’s test (P<0.05), using the statistical package SAS (Statistical Analysis System) version 9.2.

## Results

### Endobiome analysis of maize roots

When inoculated into plant cultures, PGPR can modify the endophytic microbiome and, in turn, stimulate the growth and fitness of the host. Thus, we evaluated whether the diversity and structure of the endobiome was modulated by the bioinoculation of maize plants in a monoculture system (Maize roots + UM270) and in polyculture (Maize roots + UM270 + Milpa system), using treatment of uninoculated “Maize roots ” as a control (Figure 1). The analysis was performed in triplicate using composite samples.

**Figure 1.**
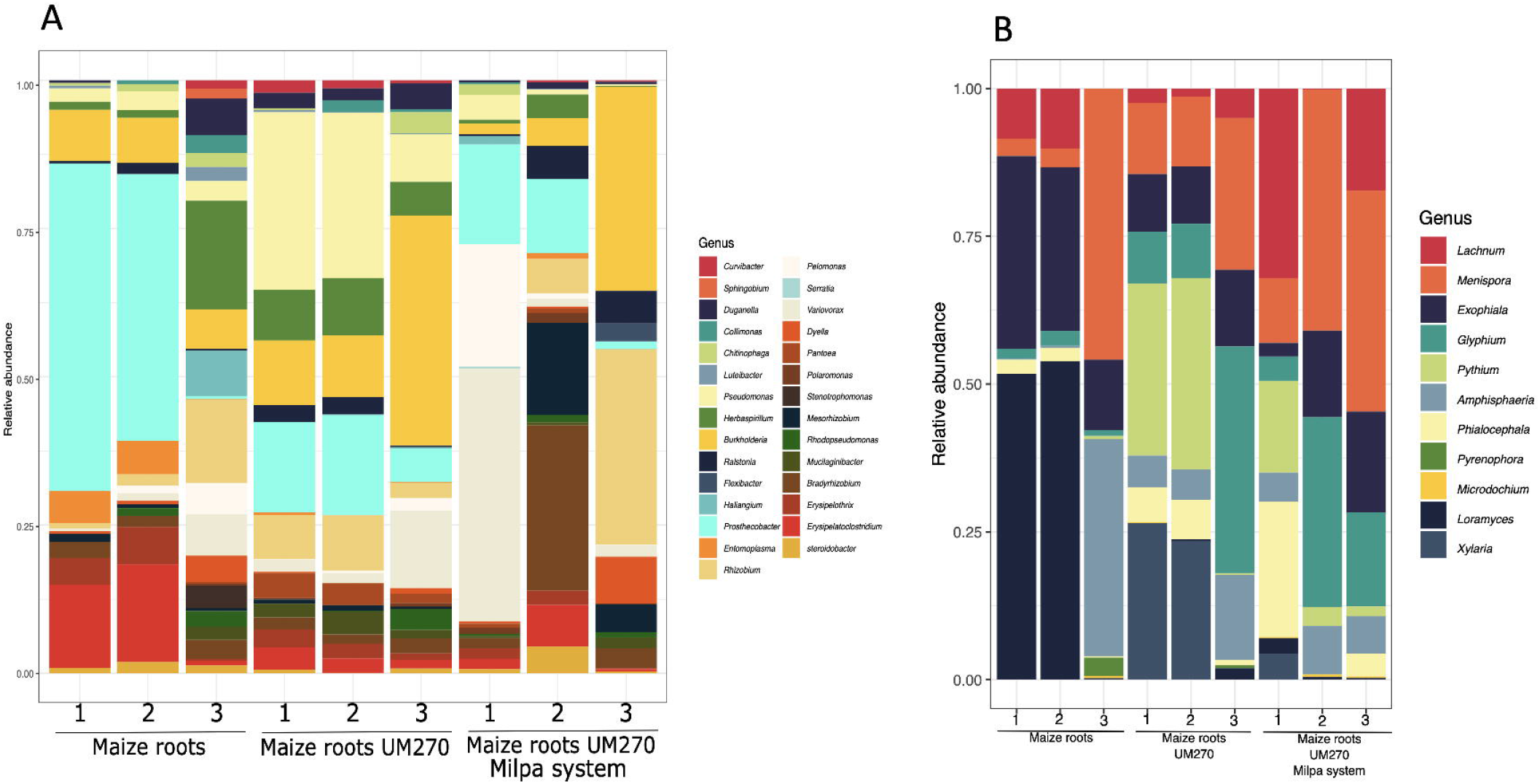
Relative abundances of bacterial (A) and fungal (B) taxa among the endophytic communities from maize plant roots cultivated in a milpa system.

The results suggested that *P. fluorescens* UM270 inoculation changed the endophytic microbiome of maize roots (Figure 1; Maize roots + UM270), compared to uninoculated plants (Figure 1; Maize roots treatment). Interestingly, Maize roots inoculated with strain UM270 showed an unexpected and very different endobiome diversity. Uninoculated maize roots highly showed abundance of OTUs belonging to genus *Prosthecobacte*r and *Curvibacter*, whose presence was decreased in inoculated treatments. On the other hand, the bacterial OTUs of the genus *Burkholderia* and *Pseudomonas* were stimulated in a monoculture (Figure 1A; Maize roots + UM270), whereas in the milpa system (Figure 1A; Maize roots + UM270 + Milpa system), the abundance of plant-associated genera, such as *Burkholderia, Variovorax*, and N-fixing rhizobia genera, such as *Rhizobium, Mesorhizobium*, and *Bradyrhizobium*, was increased.

Figure 1B shows the fungal diversity found in the maize roots from each treatment, including those biofertilized with UM270 either in mono- or polyculture (Milpa system). As noted, no significant association was correlated with the presence of the UM270 strain; however, it was interesting to detect a high abundance of mycorrhizal fungi such as *Rizophagus irregularis* or the plant growth-promoting fungus *Exophiala pisciphila*.

The increase in the number of these OTUs is better observed in Figure 2 (panels A and B) for bacterial and fungi, respectively. Some OTUs, showed in grey color, unexpectedly were increased in a milpa system, which belong to *Burkholderia* and *Variovorax* genus. Other N-fixing bacteria were also increased in inoculated plants with UM270 strain. It was also noted that some OTUs such as *Candidatus Phytoplasma* (a phytoplasma taxon associated with aster yellows disease) were also increased in one of the composed samples. However, no disease symptoms were detected in maize plants.

**Figure 2.**
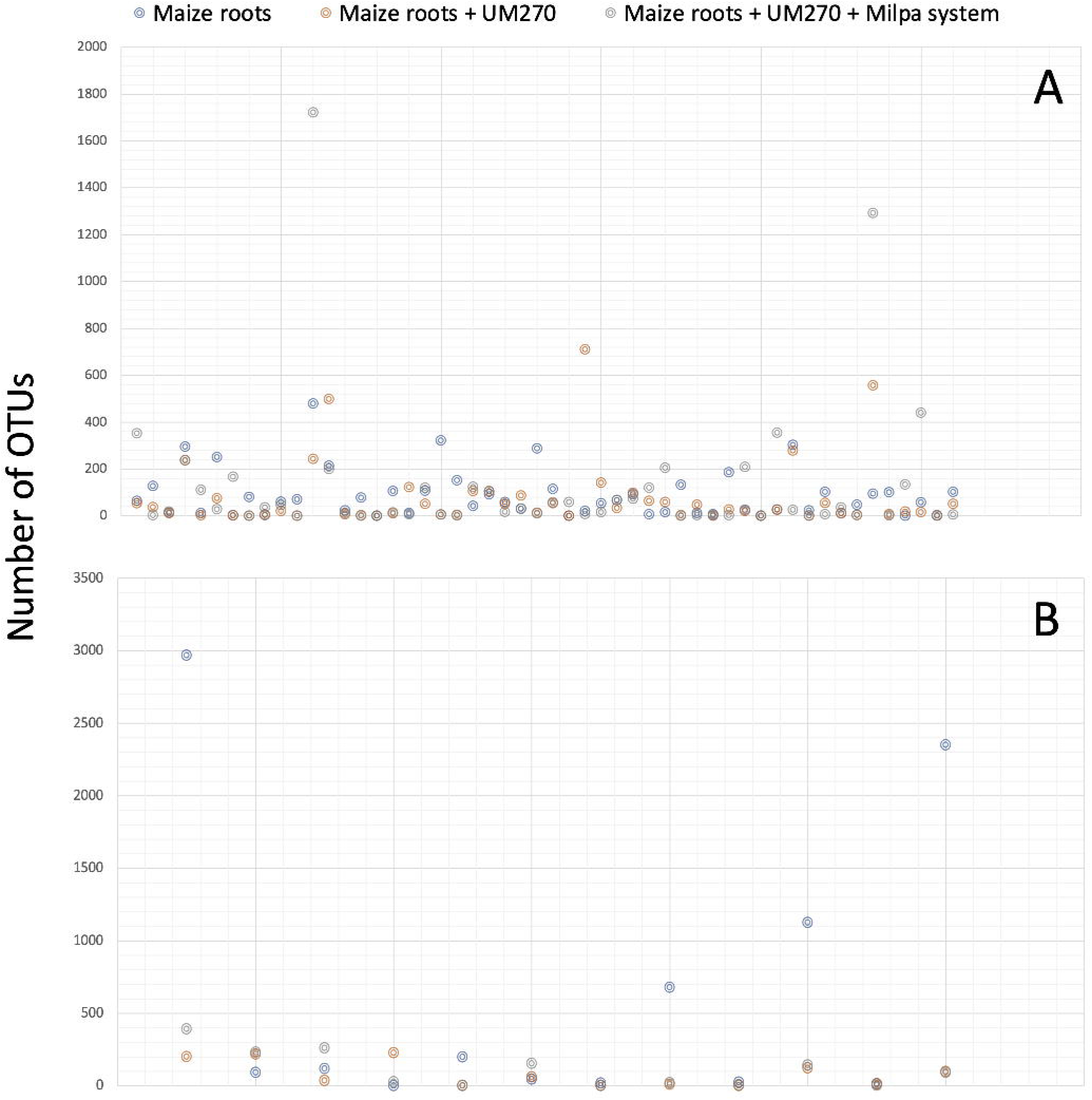
Number of OTUs detected in each of the three treatments.

In the case of the OTUs belonging to fungi, such evident results were not found where there was a correlation with the inoculation of the rhizobacterium UM270. On the contrary, it was noted that the abundance of some possible species decreased with the inoculation of the UM270 strain.

### Index diversity analysis

In this work, three of the main ecological indices were analyzed, such as Chao1, Shannon and Simpson, as shown in Figure 3A for bacteria and Figure 3B for fungi. The results show that the inoculation of the UM270 bacterium in the monoculture and polyculture treatments of corn, there is no significant change. Except in one of the treatments under the milpa system where the inoculation with the UM270 strain decreased the Alpha diversity in such maize roots. The other two treatments showed no change.

**Figure 3.**
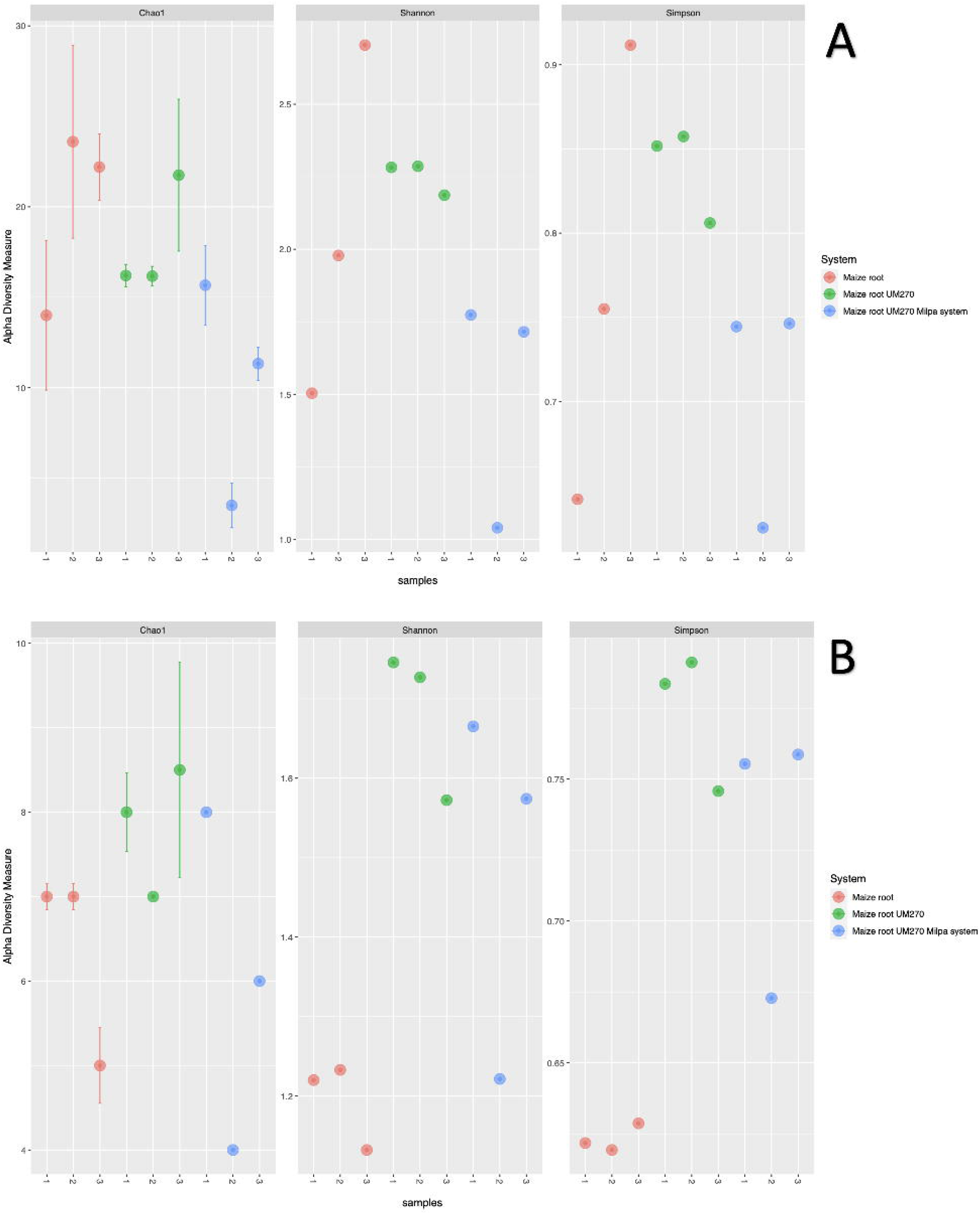
Alpha diversity indexes of bacterial (A) and fungal (B) of the endophytic communities from maize plant roots cultivated in a milpa system. A) Three replicates of bacterial and fungal diversity obtained from maize roots. B) Three replicates of bacterial and fungal diversity obtained from maize roots inoculated with UM270. C) Three replicates of bacterial and fungal diversity obtained from maize roots from a milpa system inoculated with UM270.

Figure 4 represent the shared bacterial and fungal OTUs among the three treatments. The results showed that 41 bacterial and 9 fungal OTUs were shared among the treatments; however, only 2 bacterial OTUs were found to be unique in maize roots without inoculation, and only one OUT in roots inoculated with UM270 in growing in a milpa system. No unique OTUs were found in the fungal endobiome among the treatments.

**Figure 4.**
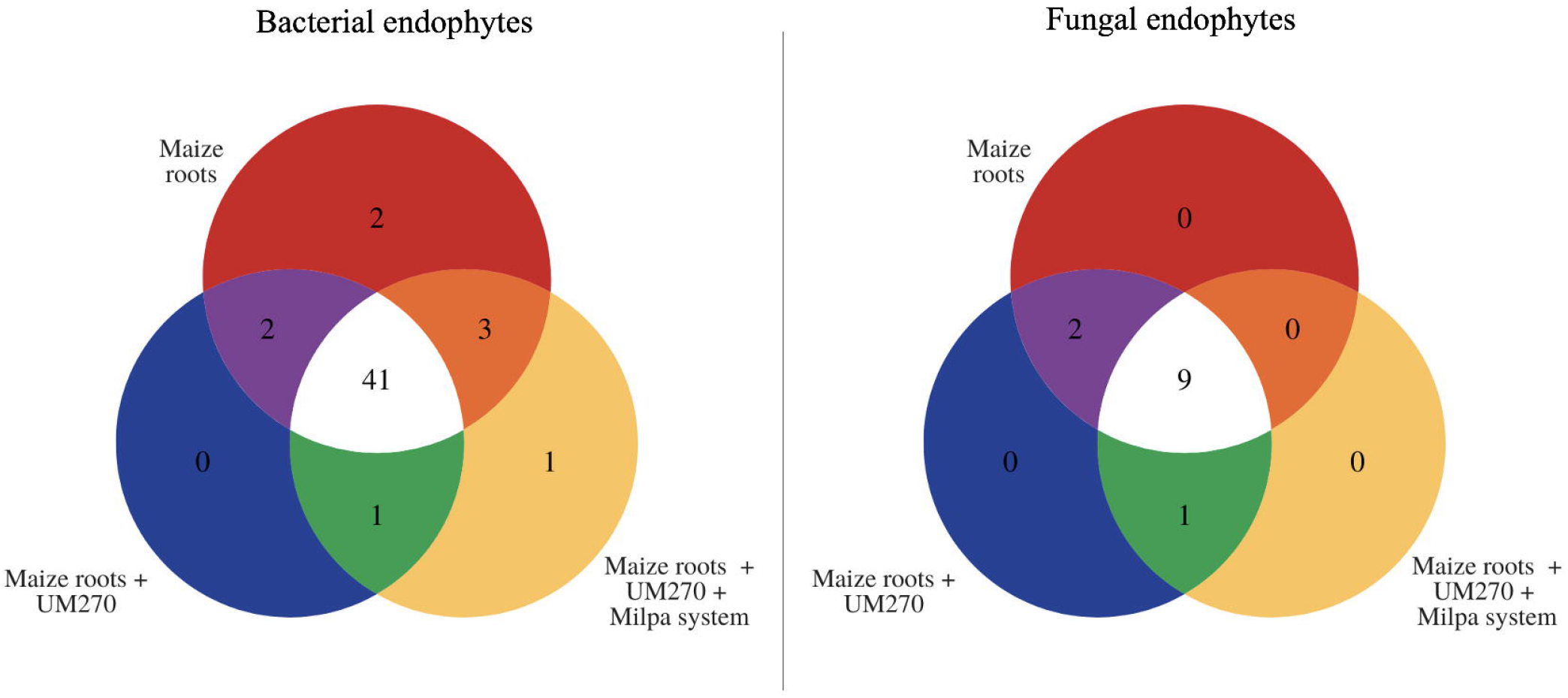
Shared OTUs among treatments.

### Endobiome Network Analysis

It was performed a network analysis to evaluate possible species interactions among the three endobiomes (Figure 5). By identifying unique, common, and co-occurring species, we can better understand the potential ecological relationships between different species and their influence on the overall health and function of endobiomes. This information can be used to develop targeted interventions to promote a healthy endobiome and prevent imbalances in the microbial communities (Trivedi et al. 2021).

**Figure 5.**
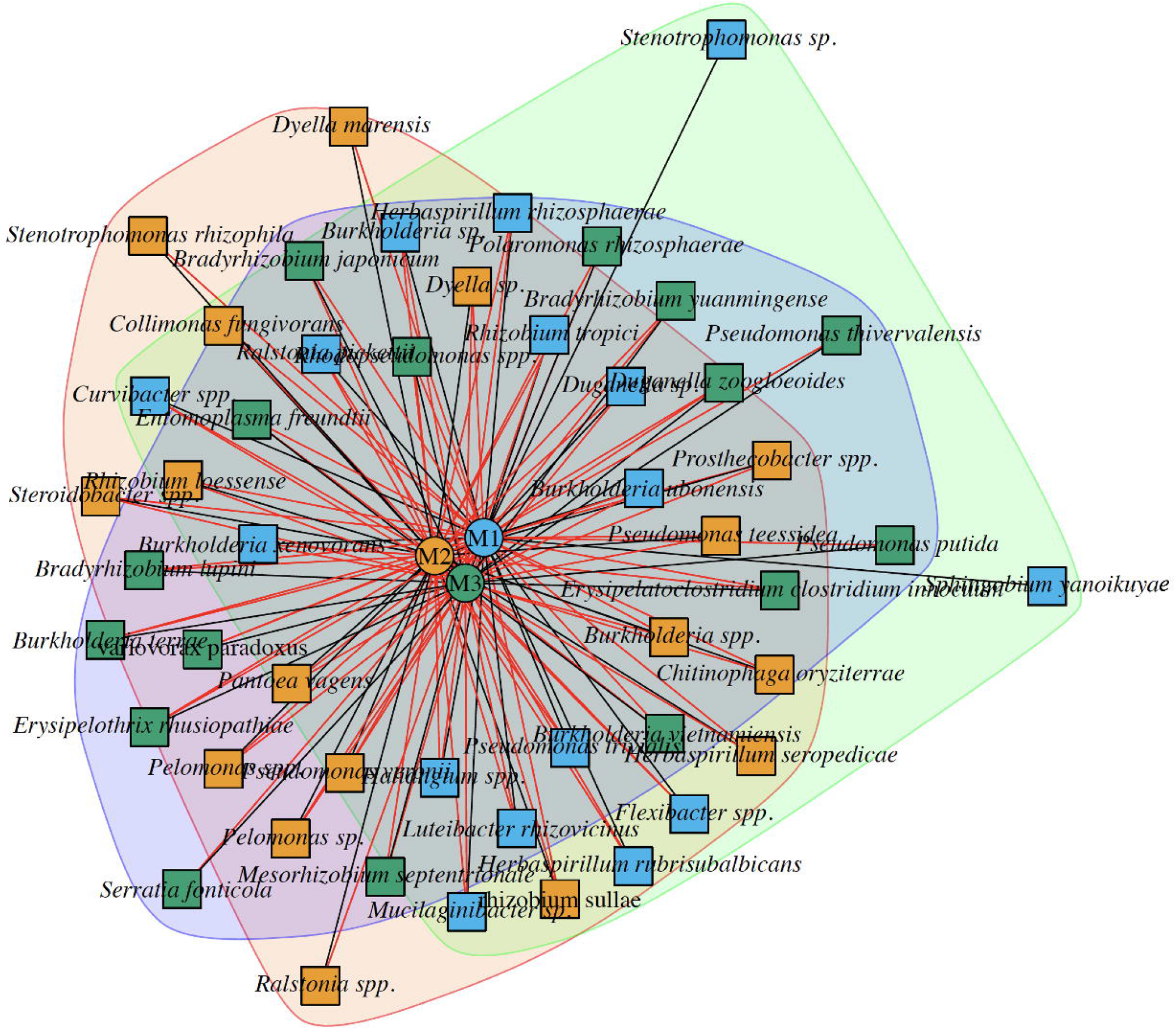
Network analysis of endophytic bacterial communities (endobiomes) from maize plant roots cultivated in a milpa system, inoculated or not with *P. fluorescens* UM270. The boxes represent individual endobiomes: M1 (maize roots), M2 (maize + root UM270), and M3 (maize + UM270 + milpa system). In the network, black lines indicate species that are unique and not present in the endobiomes. In contrast, the red lines indicate interactions or co-occurrences of species in the endobiomes.

It is interesting to note that despite the different maize treatments, multiple species were present in the endobiomes (Figure 5). This suggests that there are fundamental relationships between certain microbial species and maize plants that are not affected by specific treatments.

However, some bacterial species are found only in certain endobiomes, particularly in untreated maize. These species included *Stenotrophomonas sp*., *Burkholderia xenovorans*, and *Sphingobium yanoikuyae*. The unique occurrence of these bacteria suggests that they may play a role in corn plant growth, development, and health, especially in the absence of external treatments. For example, *Stenotrophomonas* sp. is a bacterial genus known to have a wide range of metabolic capabilities and can survive under a variety of environmental conditions (Chauviat et al. 2023). Some species of *Stenotrophomonas* have been found to be plant growth-promoting bacteria that can increase the growth and yield of crops such as maize. For example, some strains of *Stenotrophomonas* have been found to produce indole acetic acid, a plant hormone that stimulates the growth and development of maize roots.

*Burkholderia xenovorans* is another bacterial species that has been found in the endobiomes of maize plants. This species degrades various environmental pollutants, including pesticides and herbicides. This suggests that *Burkholderia xenovorans* may play a role in detoxifying the soil and protecting maize plants from the harmful effects of these chemicals (dos Santos et al. 2022).

*Sphingobium yanoikuyae* is a bacterial species known to degrade polycyclic aromatic hydrocarbons (PAHs) and environmental pollutants that are toxic to plants (Chen et al. 2021). This suggests that *Sphingobium yanoikuyae* may protect maize plants from the harmful effects of PAHs in soil. In the M2 condition (maize + root UM270), bacteria such as *Dyella marensis, Stenotrophomonas rhizophila*, and *Ralstonia* spp. co-occurred with M1 but not with M3. The co-occurrence of *Dyella marensis, Stenotrophomonas rhizophila*, and *Ralstonia* spp. with M1, but not with M3, suggests that the addition of M3 may alter the microbial community structure in the maize endobiome and instead favor the development of other microbial species. *Dyella marensis* is a bacterial species that occurs in the soil and is known for its ability to degrade a wide range of environmental pollutants. *Stenotrophomonas rhizophila* is another bacterial species known to promote plant growth and has been found in the endobiomes of several plant species, including corn. Some strains of *Stenotrophomonas rhizophila* produce plant hormones and enzymes that can stimulate root growth and plant development (Chauviat et al. 2023). Plant growth-promoting properties have also been found in some *Ralstonia* species, such as the production of plant hormones and enzymes that stimulate root growth and nutrient uptake. In the M3 system, *Pseudomonas putida, Pseudomonas thivervalensis*, and *Serratia fonticola* co-occurred only in M1 and not in M2. *Pseudomonas putida* is present in the endobiomes of several plant species, including maize, and may play a role in promoting plant growth and health (Costa-Gutierrez et al. 2022). *Pseudomonas thivervalensis* is a less well-studied bacterial species; however, some strains have been found to produce compounds that can inhibit the growth of plant pathogens (Nascimento et al. 2021). *Serratia fonticola* is a bacterial species found in various environments, including soil and water (Jung et al. 2020).

## Discussion

Corn is one of the most important cereals worldwide, and owing to its nutritional contribution, it is key to safeguarding food security, including its cultivation via the Milpa model, which is key to generating a variety of agricultural products in a small space and stimulating soil biodiversity (Gómez-Martínez et al. 2020; Hernández Galindo et al. 2022; Rodríguez and Arias de Reyna 2014).

The results of this work show that the application of a biofertilizer based on the rhizobacterium *P. fluorescens* UM270 under the milpa model modulates the microbial diversity of the root endophytes. Additionally, the inoculation of beneficial microbial agents associated with plants can engage other synergistic microbes (Santoyo 2022). Although the mechanism is not very clear, the work by Zhang et al. (2019) showed that the pre-inoculation of pepper seedlings with *Bacillus velezensis* strain NJAU-Z9 induced changes in the structure of the rhizospheric microbiome in a field experiment, stimulating communities of genera such as *Bradyrhizobium, Chitinophaga, Streptomyces, Lysobacter, Pseudomonas*, and *Rhizomicrobium* (Zhang et al. 2019). Recently, the endophytic bacteriome of Medicago truncatula was modified by the interaction of the biocompound N,N-dimethylhexadecylamine (DMHDA), produced by PGPRs such as *Arthrobacter* sp. UMCV2, and *Pseudomonas fluorescens* UM270. The results showed that bacterial groups such as β-proteobacteria and α-proteobacteria were more abundant in the root and shoot endophytic compartments, respectively (Real-Sosa et al. 2022)hidalo. Here, we observed that some genera, such as *Burkholderia, Variovorax*, and N-fixing rhizobia genera, such as *Rhizobium, Mesorhizobium* and *Bradyrhizobium*, were more abundant in maize cultures inoculated with rhizobacteria UM270. Therefore, it is also possible that these nitrogen-fixing bacteria were stimulated by nodulation factors released by bean plants, and in turn, improved the acquisition of nitrogen, one of the elements increased in the ear. It should be noted that it has been reported that intercropping between crops of faba beans (*Vicia faba* L.) and maize can result in overyielding and enhanced nodulation by faba beans (Li et al. 2016). Likewise, co-inoculation with *Rhizobium pisi* and *Pseudomonas monteilii* has been an effective biofertilization strategy for common bean production in Cuban soils (Sánchez et al. 2014). Other studies have also shown synergism between rhizobia and PGPRs to increase the growth and production of maize and beans under different environmental conditions (Coniglio et al. 2022; De Vasconcelos, Martinez Ferreira et al. 2020; Korir et al. 2017; Leite et al. 2022). However, the increase in the abundance of nitrogen-fixing genera in the endophytic (or rhizospheric) microbiome by PGPR UM270 and some mutants with low soil colonization capacity, as well as cooperation in plant nutrition, are currently being investigated.

The beneficial mycorrhizal fungus *Rizophagus irregularis* was detected in maize roots in the three treatments analyzed, including the milpa system. Some previous studies shows that *R. irregularis* can promote the growth of bean plants under greenhouse conditions, as well as under field conditions, having positive effects on maize, soybean and wheat (Hidalgo Rodríguez et al. 2019; Renaut et al. 2020). Similarly, the endophytic fungus *Exophiala pisciphila*, particularly the H93 strain, has been an excellent promoter of plant growth in maize, particularly under conditions of stress caused by heavy metals. One action of *E. pisciphila* is to improve plant nutrition by solubilizing phosphates (Xu *et al*. 2020). Other species found as endophytes of maize were *Menispora tortuosa, Glyphium elatum* or *Phialocephala subalpina*, to mention a few, but they have been more associated with woody plants (Lorenzo and Messuti 2005; Réblová et al. 2006; Schlegel et al. 2016); however, it would be interesting to explore its symbiotic functions with plants of agricultural interest.

Finally, Beta diversity detected in the endophytic microbiome of maize roots in monoculture and biofertilized with *P. fluorescens* UM270 showed the lowest biodiversity variations with respect to the other treatments. Although it can be argued that polyculture (or cropping practices) and fertilization with biological agents can stimulate greater endophytic diversity (Srivastava et al. 2007), the uninoculated maize monoculture also showed high variation.

Finally, the endobiome network allowed the identification of different bacterial species present in the three treatment types of maize, indicating the presence of fundamental relationships between certain microbial species and maize plants that were not affected by the specific treatments. In addition, some unique bacterial species were identified in specific endobiomes (e.g. *Stenotrophomonas* sp. or *Burkholderia* sp.), indicating their possible roles in the growth, development, and health of corn plants, especially in the absence of external treatments.

## Conclusion

The addition of biofertilizers in maize plants growing under milpa models, such as the *P. fluorescens* UM270 strain, modulates the endophytic microbiome. One of the potential mechanisms employed by the UM270 strain to stimulate plant growth might the recruitment of other beneficial endophytic microorganisms. However, this hypothesis requires further investigation trough isolation and characterization of synergistic activities of the inoculating strain UM270 and the resident endophytic organisms on maize plants.

## Acknowledgements

G.S. thanks CONACYT-México (Proposal: A1-S-15956) and CIC-UMSNH (2021-2022) for financial support to our research projects.

